# Nonlinear decoding of natural images from large-scale primate retinal ganglion recordings

**DOI:** 10.1101/2020.09.07.285742

**Authors:** Young Joon Kim, Nora Brackbill, Ella Batty, JinHyung Lee, Catalin Mitelut, William Tong, E.J. Chichilnisky, Liam Paninski

## Abstract

Decoding sensory stimuli from neural activity can provide insight into how the nervous system might interpret the physical environment, and facilitates the development of brain-machine interfaces. Nevertheless, the neural decoding problem remains a significant open challenge. Here, we present an efficient nonlinear decoding approach for inferring natural scene stimuli from the spiking activities of retinal ganglion cells (RGCs). Our approach uses neural networks to improve upon existing decoders in both accuracy and scalability. Trained and validated on real retinal spike data from > 1000 simultaneously recorded macaque RGC units, the decoder demonstrates the necessity of nonlinear computations for accurate decoding of the fine structures of visual stimuli. Specifically, high-pass spatial features of natural images can only be decoded using nonlinear techniques, while low-pass features can be extracted equally well by linear and nonlinear methods. Together, these results advance the state of the art in decoding natural stimuli from large populations of neurons.

**Author summary:** Neural decoding is a fundamental problem in computational and statistical neuroscience. There is an enormous literature on this problem, applied to a wide variety of brain areas and nervous systems. Here we focus on the problem of decoding visual information from the retina. The bulk of previous work here has focused on simple linear decoders, applied to modest numbers of simultaneously recorded cells, to decode artificial stimuli. In contrast, here we develop a scalable nonlinear decoding method to decode natural images from the responses of over a thousand simultaneously recorded units, and show that this decoder significantly improves on the state of the art.

## Introduction

What is the relationship between stimuli and neural activity? While this critical neural coding problem has often been approached from the perspective of developing and testing encoding models, the inverse task of *decoding* — the mapping from neural signals to stimuli — can provide insight into understanding neural coding. Furthermore, efficient decoding is crucial for the development of brain-computer interfaces and neuroprosthetic devices [1–10].

The retina has long provided a useful testbed for decoding methods, since mapping retinal ganglion cell (RGC) responses into a decoded image provides a direct visualization of decoding model performance. Most approaches to decoding images from RGCs have depended on linear methods, due to their interpretability and computational efficiency [1, 11, 12]. Although linear methods successfully decoded spatially uniform white noise stimuli [1] and the coarse structure of natural scene stimuli from RGC population responses [12], they largely fail to recover finer visual details of naturalistic images.

More recent decoders incorporate nonlinear methods for more accurate decoding of complex visual stimuli. Some have leveraged optimal Bayesian decoding for white noise stimuli, but exhibited limited scalability to large neural populations [13]. Others have attempted to incorporate key prior information for natural scene image structures and perform computationally expensive approximations to Bayesian inference [14, 15]. Unfortunately, computational complexity and difficulties in formulating an accurate prior for natural scenery have hindered these methods. Other studies have constructed decoders that explicitly model the correlations between spike trains of different cells, for example by using the relative timings of first spikes as the measure of neural response [16]. Parallel endeavors into decoding calcium imaging recordings from the visual cortex have produced coarse reconstructions of naturalistic stimuli through both linear and nonlinear approaches [17–20].

In parallel, some recent decoders have relied on neural networks as efficient Bayesian inference approximators. However, established neural network decoders have either only been validated on artificial spike datasets [21–23] or on limited real-world datasets with modest numbers of simultaneously recorded cells [23–25]. No nonlinear decoder has been developed and evaluated with the ultimate goal of efficiently decoding natural scenes from large populations (e.g., thousands) of neurons. Because the crux of the neural coding problem is to understand how the brain encodes and decodes naturalistic stimuli in through large neuronal populations, it is crucial to address this gap.

Therefore in this work we developed a multi-stage decoding approach that exhibits improved accuracy over linear methods and greater efficiency over existing nonlinear methods, and applied this decoder to decode natural images from large-scale multi-electrode recordings from the primate retina.

## Results

### Overview

All decoding results were obtained on retinal datasets consisting of macaque RGC spike responses to natural scene images [12]. Two identically prepared datasets, each containing responses to 10,000 images, were used for independent validation of our decoding methods. The electrophysiological recordings were spike sorted using YASS [26] to identify 2094 and 1897 natural scene RGC units for the two datasets. We also recorded the responses to white noise visual stimulation and estimated receptive fields to classify these units into retinal ganglion cell types, to allow for analyses of cell-type specific natural scene decoding. See **Materials and methods** for full details.

Our decoding approach addresses accuracy and scalability by segmenting the decoding task into three sub-tasks (**Figure 1**).

- We use linear ridge regression to map the spike-sorted, time-binned RGC spikes to “low-pass,” Gaussian-smoothed versions of the target images. The smoothing filter size approximates the receptive fields of ON and OFF midget RGCs, the cell types with the highest densities in the primate retina.
- A spatially-restricted neural network decoder is trained to capture the nonlinear relationship between the RGC spikes and the “high-pass” images, which are the residuals between the true and the low-pass images from the first step. The high-pass and low-pass outputs are summed to produce “combined” decoded image s (**Figure 2**).
- A deblurring network is trained and applied to improve the combined decoder outputs by enforcing natural image priors.

**Fig. 1.**
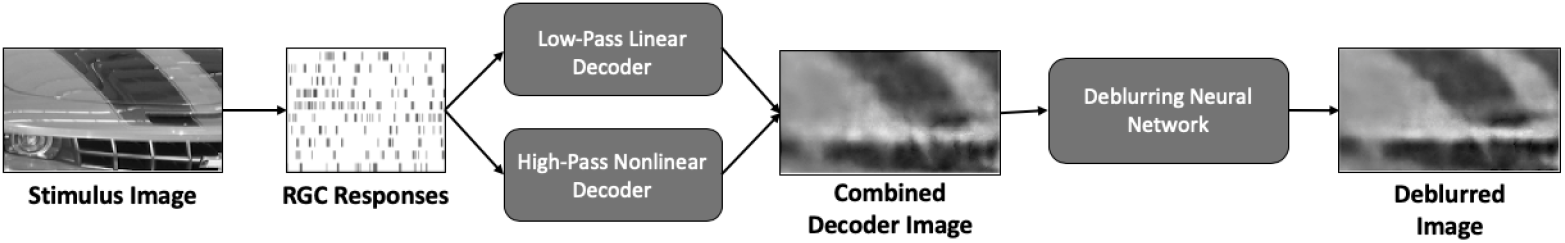
Outline of the decoding method. RGC responses to image stimuli are passed through both linear and nonlinear decoders to decode the low-pass and high-pass components of the original stimuli, respectively, before the combined decoded images are deblurred and denoised by a separate deblurring neural network.

The division of visual decoding into low-pass and high-pass decoding sub-tasks allowed us to leverage linear regression, which is simple and quick, for obtaining the target images’ global features, while having the neural network decoder focus its statistical power on the addition of finer visual details. As discussed below, this strategy yielded better results than applying the neural network decoder to either the low-pass or the whole test images (**Table 1**).

**Table 1.**
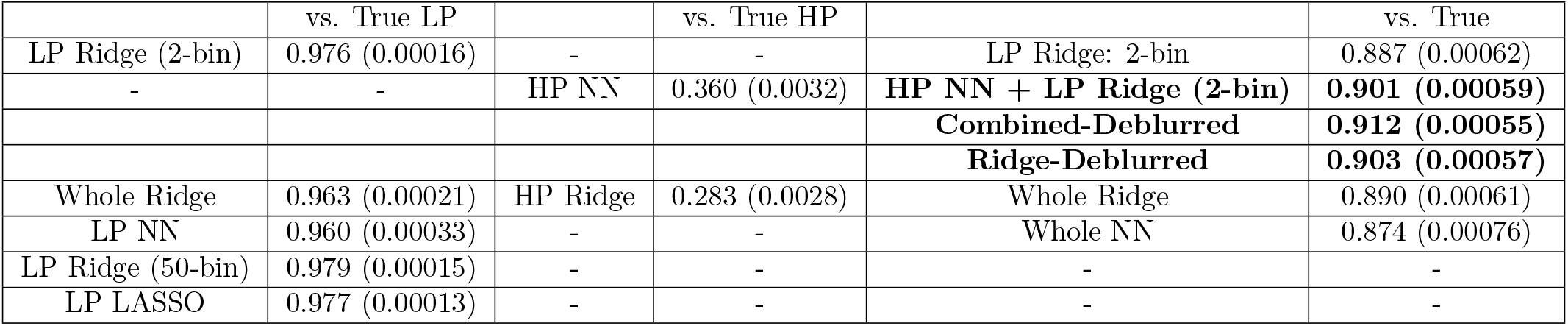
Pixel-wise test correlations of all decoder outputs (99% confidence interval values in parentheses). The 2-bin and 50-bin LP ridge labels represent the two linear ridge decoders trained on the low-pass images. The whole ridge decoder is the 2-bin ridge decoder trained on the true whole images themselves while the HP ridge decoder is the same decoder trained on the high-pass images only. The LP, HP, and whole NN labels denote the spatially restricted neural network decoder trained on low-pass, high-pass, and whole images, respectively. LP LASSO represents the 2-bin LASSO regression decoder trained on low-pass images. Finally, the combined-deblurred images are the deblurred versions of the sum of the HP NN and LP Ridge (2-bin) decoded images while the ridge-deblurred images are the deblurred versions of the whole ridge decoder outputs. These final three (combined-deblurred, ridge-deblurred, and HP NN + LP Ridge (2-bin)) are bolded as they produced best results. The second, fourth, and sixth columns represent pixel-wise test correlations of each decoder’s output versus the true low-pass, high-pass, and whole images, respectively.

### Linear decoding efficiently decodes low-pass spatial features

We used two penalized linear regression approaches (ridge and LASSO regression [27]) for linearly decoding the low-pass images. Both decoders only considered the neural responses during the image onset (30 - 150 ms) and offset (170 - 300 ms) time frames (**Figure 3, Top Left**). For reference, LASSO regression is a form of linear regression whose regularization method enforces sparsity such that the uninformative input variables are assigned zero weights while the informative inputs are assigned non-zero weights [27]. In the process, LASSO successfully identified each RGC unit’s relevant linear spatial weights for both the image onset and offset time bins while zero-ing out the insignificant spatial weights (**Figure 3, Bottom**).

**Fig. 3.**
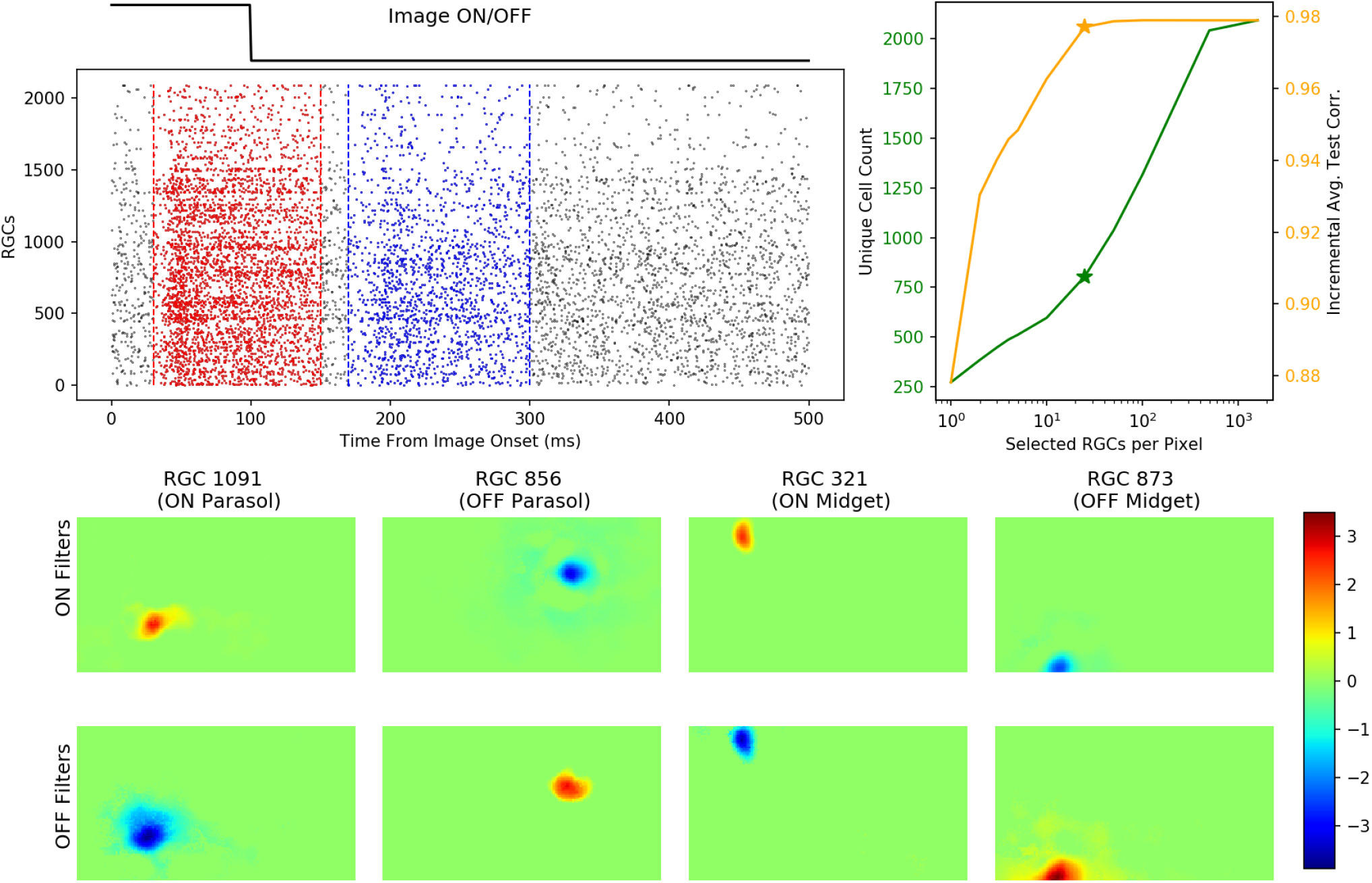
LASSO regression establishes a sparse mapping between RGC units and pixels. (**Top Left**) Schematic of the ON (red; 30 - 150 ms) and OFF (blue; 170 - 300 ms) responses derived from RGC spikes. Each RGC’s ON and OFF filter weights were multiplied to the summed spike counts within these windows. The spikes in these bins represent the cells’ responses to stimuli onsets and offsets, respectively. The raster density (each dot represents a spike from a single RGC unit on a single trial) indicates that most of the RGC units’ spikes were found in these two bins, which came slightly after the stimuli onsets and offsets themselves as shown by the top line. (**Top Right**) Total unique selected RGC unit count (green) and mean pixel-wise test correlations of partial LASSO decoded images (orange) as functions of the number of units chosen per pixel. For each pixel, {1, 2, 3, 4, 5, 10, 25, 50, 100, 500, 1000, 1600} top units were chosen. Asterisks mark top 25 units per pixel (805 unique units and 0.977 test correlation), the hyperparameter setting chosen for the nonlinear decoder below. (**Bottom**) Representative “ON” and “OFF” spatial weights estimated by LASSO regression for four RGC units. Overall, LASSO regression successfully established a sparse mapping between RGC units and individual pixels by zeroing each cells’ uninformative spatial weights, which comprise the majorities of the ON and OFF filters.

The LASSO spatial filters were roughly similar in appearance to the corresponding RGC unit receptive fields calculated from spike-triggered averages of white noise recordings (data not shown; see [12]). These linear filters eventually allowed for a sparse mapping between RGC units and image pixels so that only the most informative units for each pixel would be used as inputs for the nonlinear decoder [25]. Partial LASSO-based decoding using smaller subsets of informative units demonstrated that these few hundred units were responsible for most of the decoding accuracy observed (**Figure 3, Top Right**). Ultimately, 25 top units per pixel, corresponding to 805 total unique RGC units and a mean low-pass test correlation of 0.977 (±0.0002; this and all following error bars correspond to 99% CI values), were chosen. Choosing fewer than 25 informative RGC units per pixel resulted in lower LASSO regression test correlations, while choosing more units per pixel increased computational load without concomitant improvements in test correlation.

Consistent with previous findings [12], both linear decoders successfully decoded the global features of the stimuli by accurately modeling the low-pass images (**Figure 4**). When evaluated by mean pixel-wise correlation against the true low-pass images, the decoded outputs from the ridge and LASSO decoders registered test correlations of 0.976 (±0.0002) and 0.977 (±0.0001), respectively (**Figure 4**; **Table 1**)^1^. Increasing the temporal resolution of linear decoding beyond the two onset and offset time bins did not yield significant improvements in accuracy.

**Fig. 4.**
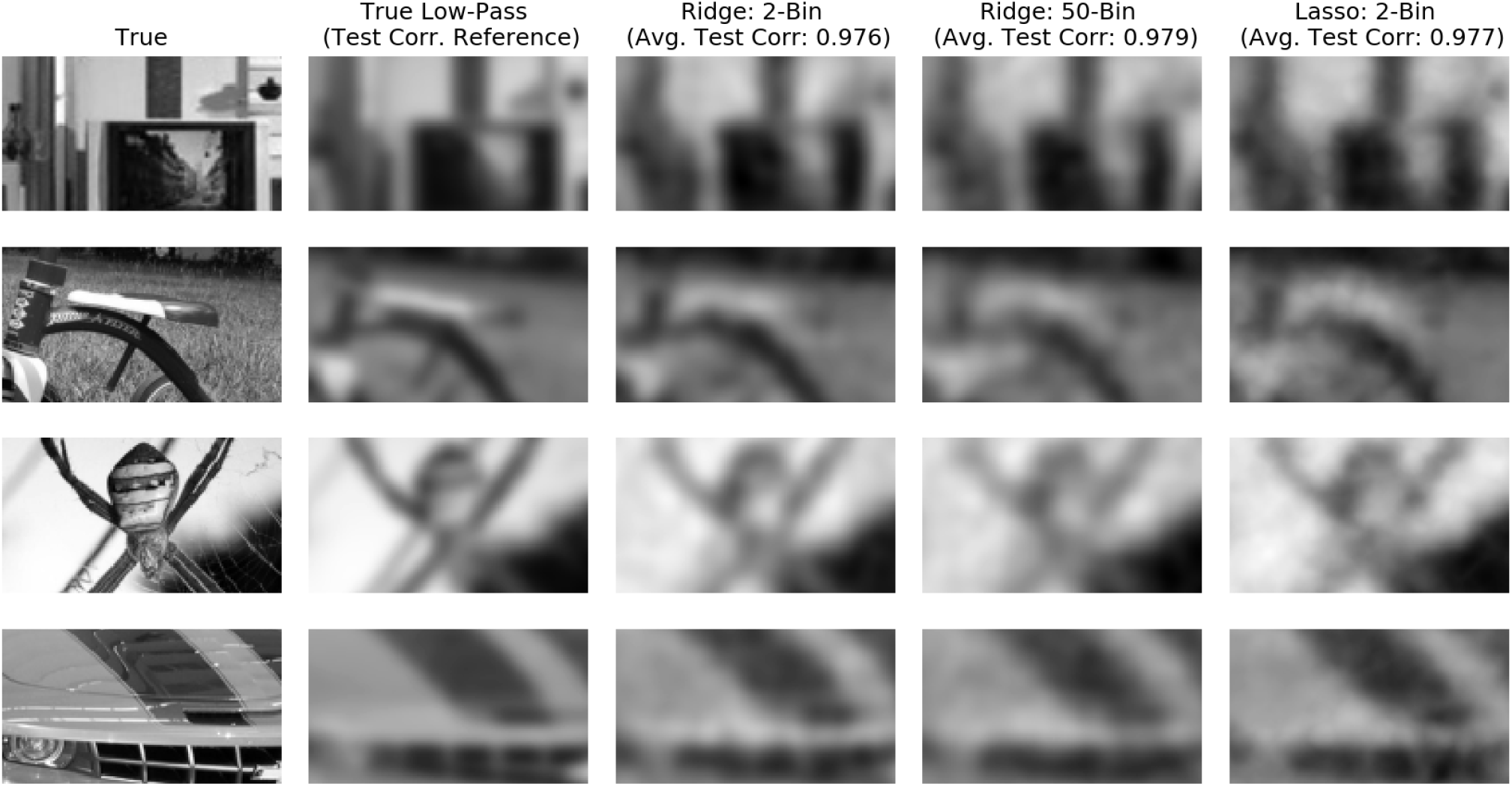
Linear decoding efficiently decodes low-pass spatial features. Representative true and true low-pass images along with their decoded low-pass counterparts produced via ridge (2-time-bin and 50-time-bin) and LASSO regression. Mean pixel-wise test correlation (evaluated against the true low-pass images, not the true images) are indicated within the top labels. The 50-bin decoder considers spike counts from the entire 500 ms stimulus window organized into 10ms bins; this decoder achieved similar accuracy as the 2-bin decoder. All three linear regression techniques produce highly accurate decoding of the true low-pass images, suggesting that linear methods are sufficient for extracting the global features of natural scene image stimuli.

How different are decoding results if the linear decoder is, instead, applied to the true whole images rather than the low-pass images, or if a nonlinear decoder is used for the low-pass targets? Notably, a ridge regression decoder trained on true images exhibited performance no better than the low-pass-specific linear decoders. Specifically, it registered a test correlation of 0.963 (±0.0002) versus true low-pass images and 0.890 (±0.0006) versus true images, suggesting that linear decoding can only recover low-pass details regardless of whether the decoding target contains high-pass details or not (**Table 1**). The ridge low-pass decoded images registered a test correlation of 0.887 (±0.0006) against the whole test images. On the other hand, applying our neural network decoder to the low-pass targets demonstrates that linear decoding is slightly more accurate (likely due to slight overfitting by the neural network) and vastly more efficient for low-pass decoding as the former exhibited a lower test correlation of 0.960 (±0.0003) versus the low-pass targets (**Table 1**). In sum, linear decoding is both the most accurate and appropriate approach for extracting the global features of natural scenes.

### Nonlinear methods improve decoding of high-pass details and utilize spike temporal correlations

Despite the high accuracy of low-pass linear decoding, the low-pass images and their decoded counterparts are (by construction) lacking the finer spatial details of the original stimuli. Therefore we turned our attention next to decoding the spatially high-pass images formed as the differences of the low-pass and original images. Again we compared linear and nonlinear decoders; unlike in the low-pass setting, we found that nonlinear decoders were able to extract significantly more information about the high-pass images than linear decoders. Specifically, a neural network decoder that used the non-zero LASSO regression weights to select its inputs (**Figure 2, Top Right**) achieved a test correlation of 0.360 (±0.003) when evaluated against the high-pass stimuli, compared to ridge regression’s test correlation of 0.283 (±0.003) (**Figure 5, Bottom Left**).

**Fig. 2.**
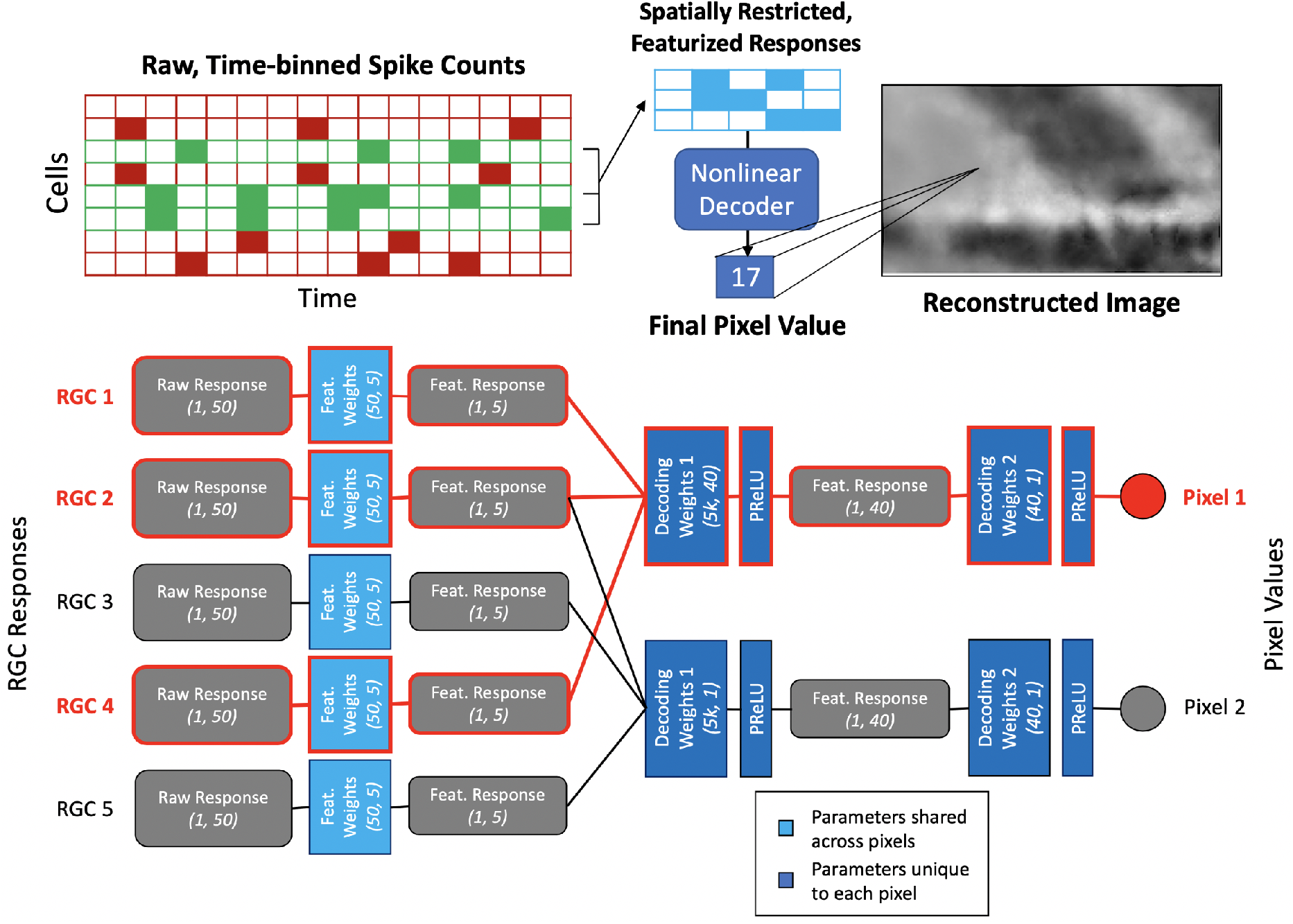
Outline of the nonlinear decoder. (**Top**) The first part of the nonlinear decoder featurizes the RGC units’ time-binned spike responses (50-dimensional vector for each RGC) to a lower dimension (*f* = 5). Afterwards, each pixel’s *k* = 25 most relevant units’ featurized vectors are gathered and passed through a spatially restricted neural network where each pixel is assigned its own nonlinear decoder to produce the final pixel value. (**Bottom**) A miniaturized schematic of the spatially restricted neural network. Parameters that are shared across pixels versus those that are unique to each pixel are color coded in different shades of blue. Furthermore, all the input values and weights that feed into a single pixel value are outlined in red to indicate the spatially restricted nature of the network. The vector dimensions of the weights and inputs are written in italicized parentheses; *k* represents the number of top units per pixel chosen for decoding.

Moreover, the combined decoder output (summing the linearly decoded low-pass and nonlinearly decoded high-pass images) consistently produced higher test correlations compared to a simple linear decoder. Relative to the true images, ridge regression (for the whole images) and combined decoding yielded mean correlations of 0.890 (±0.0006) and 0.901 (±0.0006), respectively (**Figure 5, Top**). In comparison, the linear low-pass decoded images alone yielded 0.887 (±0.0006). In other words, linear decoding of the whole image is almost no better than simply aiming for the low-pass image and nonlinear decoding is necessary to recover significantly more detail beyond the low-pass target. Additionally, a neural network decoder that targets the whole true images falls short of the combined decoder with a mean test correlation of 0.874 (±0.0008) versus true images (**Table 1**). In conjunction with the previous section’s finding that the neural network decoder is not as successful with low-pass decoding as linear decoders, these results further justify our approach to reserve nonlinear decoding for the high-pass and linear decoding for the low-pass targets.

**Fig. 5.**
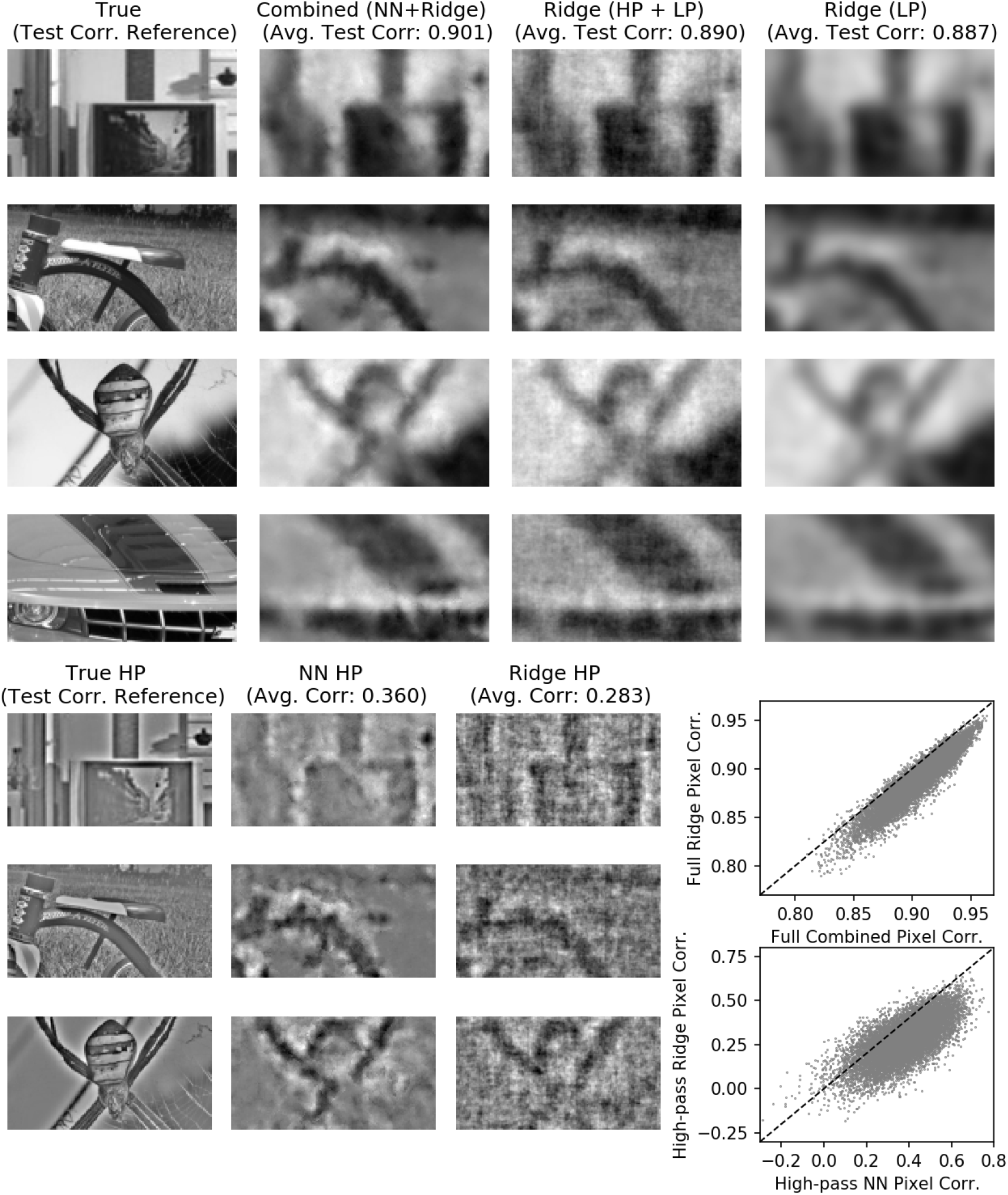
Nonlinear decoding extracts high-pass features more accurately than linear decoding. (**Top**) Representative true images with their linearly decoded and combined decoder outputs; note that the linear decoder here decodes the true images (not just the true low-pass images) and was included for overall comparison. The correlation values here compare the aforementioned decoded outputs against the true images. (**Bottom Left**) Representative high-pass images with corresponding nonlinear and linear decoded versions. The correlation values here compare the high-pass decoded outputs against the true high-pass images. (**Bottom Right**) Pixel-wise test correlation comparisons of linear and nonlinear decoding performance for the true and high-pass images. Linear decoding, either for the whole or low-pass images, is distinctly insufficient and nonlinear methods are necessary for accurate decoding.

We then sought to analyze what characteristics of the RGC spike responses allowed for the superior performance of the combined decoding method. Previous studies have reported that nonlinear decoding better incorporates spike train temporal structure, which leads to its improvement over linear methods [25, 28, 29]. However, these studies were conducted with simplified random or white noise stimuli and it is unclear how these findings translate to natural scene decoding. Thus, we hoped to shed light on how spike train correlations, both cross-neuronal and temporal, contribute to linear and nonlinear decoding. In previous literature, the former have been referred to as “noise correlations” and the latter as “history correlations” [25]. On a separate dataset of 150 test images each repeated 10 times, we created modified neural responses with either each unit’s full response being shuffled between repeat trials (removing cross-neuronal correlations) or the spike counts for each time bin being shuffled (removing temporal correlations; **Figure 6, Top**). Both transformations do not change the average firing rate over time associated with each RGC.

**Fig. 6.**
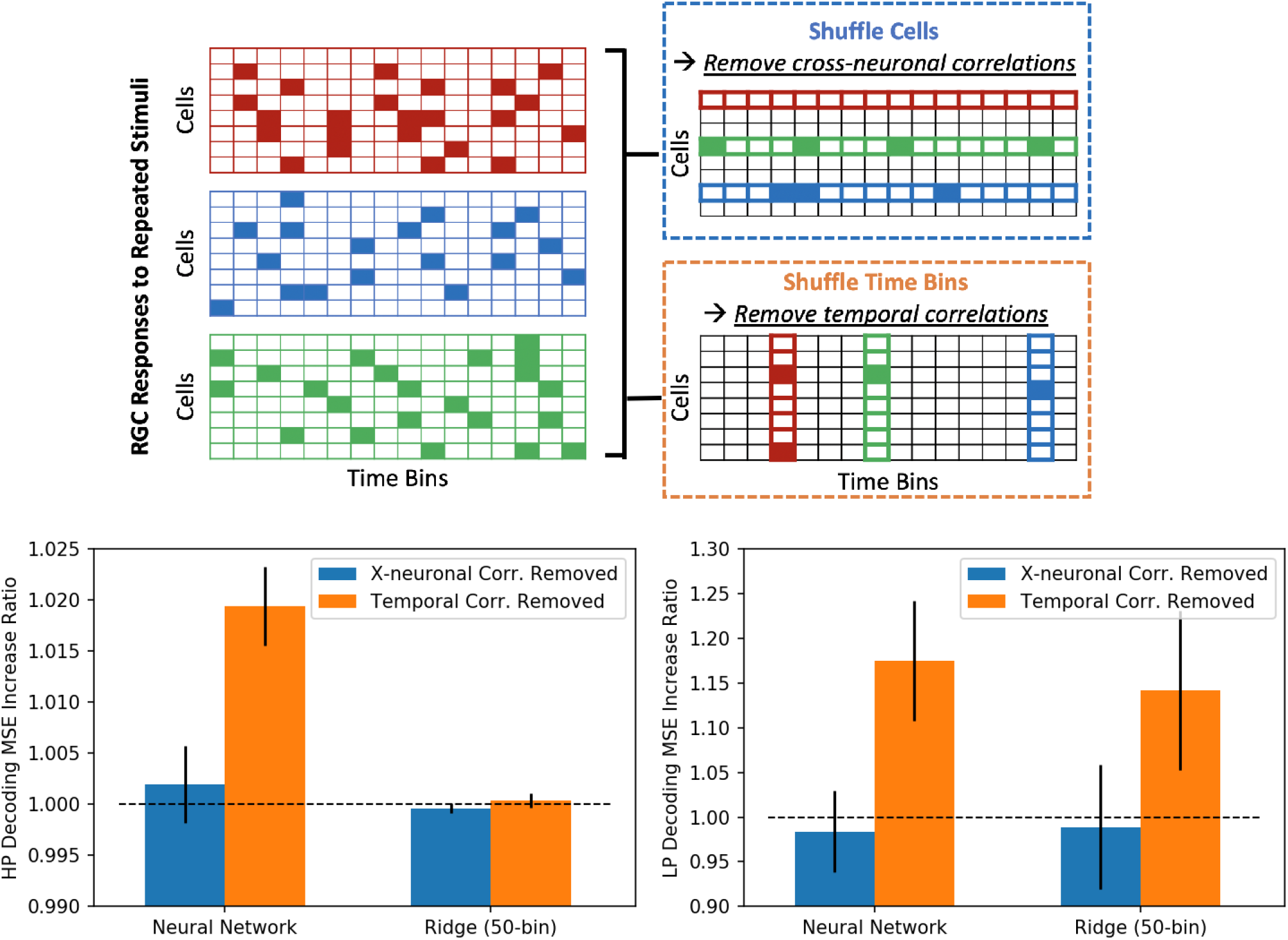
Spike temporal correlations are useful for high-pass nonlinear decoding and for low-pass decoding. (**Top**) Schematic of the shuffling of time bins and units’ responses across repeated stimuli trials. (**Bottom**) Ratio increases in MSE for neural network and linear decoders for high-pass and low-pass images before and after removing spike train correlations. While temporal correlations are important for both decoders in low-pass decoding, only the neural network decoder is reliant on temporal correlations in high-pass decoding. Cross-neuronal correlations are not crucial for both decoders in either decoding scheme.

For high-pass decoding, the neural network decoder exhibited a 1.9% (±0.4) increase in pixel-wise MSE when temporal correlations were removed, while the ridge decoder experienced a 0.04% (±0.07) increase in MSE (**Figure 6, Bottom**); i.e., nonlinear high-pass decoding is dependent on temporal correlations while linear high-pass decoding is not. Removing cross-neuronal correlations yielded no significant changes in either decoder, consistent with [12]. Meanwhile, for low-pass decoding, both decoders were equally and significantly affected by removing temporal correlations, as indicated by the 17.5% (±6.7) and 14.2% (±8.9) increases in MSE for the neural network and linear decoders, respectively (**Figure 6, Bottom**). For the above comparisons, the ridge linear decoder for 50 time bins was used to maintain the same temporal resolution as the neural network decoder. In short, spike temporal correlations are important, specifically for the low-pass linear and all nonlinear decoders for optimal performance, while cross-neuronal correlations are not influential in any decoding setup analyzed here.

### OFF midget RGC units drive improvements in high-pass decoding when using nonlinear methods

Next, we sought to investigate the differential contributions of each major RGC type towards visual decoding. Previous work has revealed that, in the context of linear decoding, midget cells convey more high frequency visual information while parasol cells tend to encode more low frequency information, consistent with the differences in density and receptive field size of these cell classes [12]. Here we focused on the ON/OFF parasol/midget cells, the four numerically dominant RGC types, and their roles in linear versus nonlinear decoding. We classified the RGCs recorded during natural scene stimulation by first identifying units recorded during white noise stimulation and then using a conservative matching scheme that ensured one-to-one matching between recorded units in the two conditions. In total, 1033 units were matched, within which there were 72 ON parasol, 87 OFF parasol, 175 ON midget, and 195 OFF midget units (**Materials and Methods**).

We performed standard ridge regression decoding for whole and low-pass images using spikes from the above four cell types and compared these decoded outputs to those derived from all 2094 RGC units, which include those not belonging to the four main types (**Figure 7**). Consistent with previous results [12], midget decoding recovers more high frequency visual information than parasol decoding, while ON and OFF units yield decoded images of similar quality. Meanwhile, differences between parasol and midget cell decoding are reduced for low-pass filtered images, as this task is not asking either cell population to decode high frequency visual information.

**Fig. 7.**
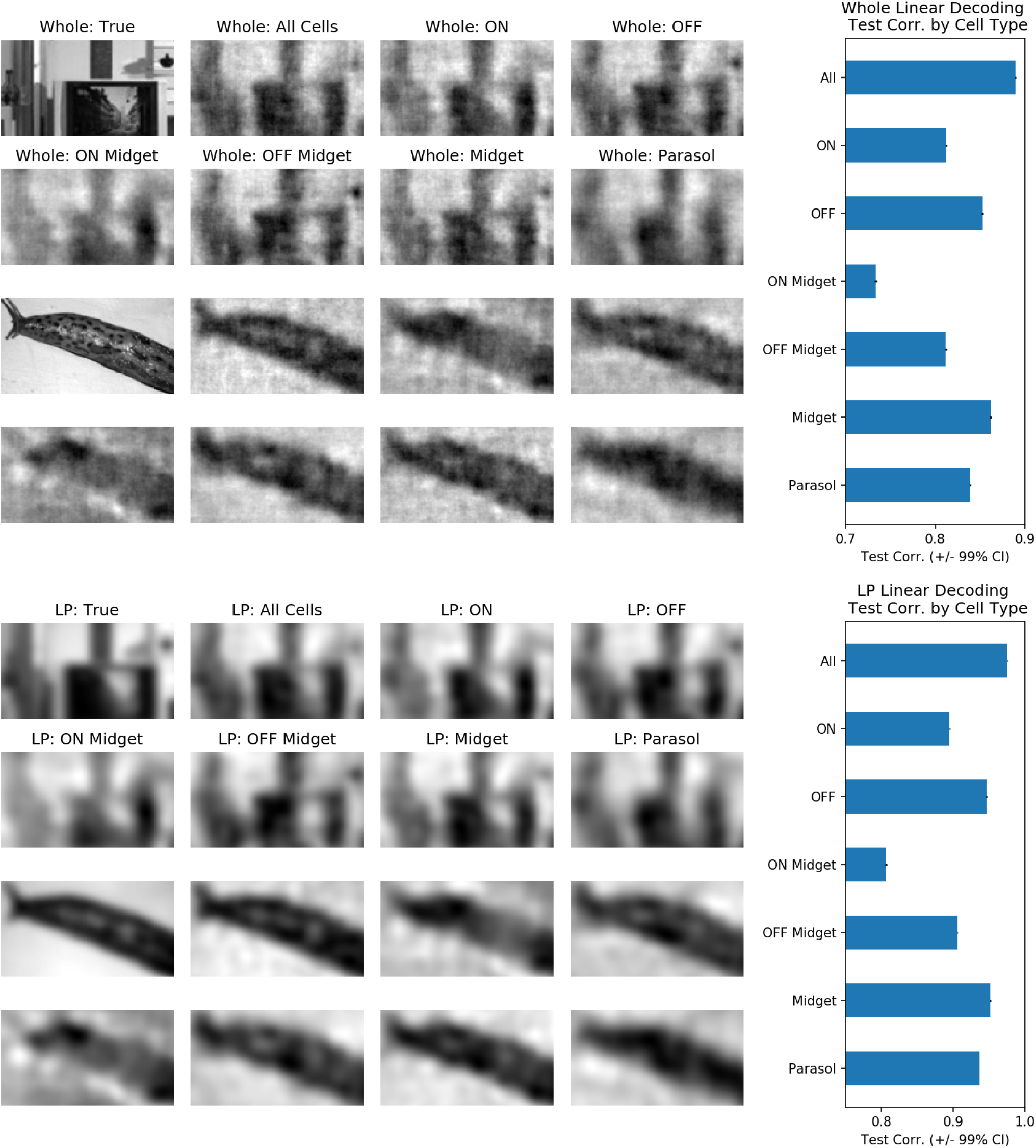
All major RGC types meaningfully contribute to low-pass linear decoding. (**Top Left**) Representative whole images with their corresponding linearly decoded outputs using all, ON, OFF, ON Midget, OFF midget, midget, and parasol units, respectively. (**Top Right**) Whole test correlations as functions of RGC type used for linear decoding. (**Bottom Left**) Representative low-pass images with their corresponding linearly decoded outputs using all, ON, OFF, ON Midget, OFF midget, midget, and parasol units, respectively. (**Bottom Right**) Low-pass test correlations as functions of RGC type used for linear decoding. Midget units encode more high frequency information than parasol units while ON and OFF units produce similar qualities of decoding. Overall, all RGC types contribute meaningfully to low-pass, linear decoding.

We then investigated cell type contributions in the context of high-pass decoding (**Figure 8**). Specifically, we investigated which cell type contributed most to the advantage of nonlinear over linear high-pass decoding and, thus, explained the improved performance of our decoding scheme. The advantages of nonlinear decoding were most prominent for midget and OFF units, with mean increases in test correlation by 7.1% and 6.8%, respectively (**Figure 8, Top Right**). Parasol and ON units, meanwhile, saw a statistically insignificant change in test correlation. More fine-grained analyses showed that only the OFF midget units enjoyed a statistically significant increase of 6.5% in mean test correlation in high-pass decoding. While ON midget units did indeed contribute meaningfully to high-pass decoding (as shown by their relatively high test correlations), they enjoyed no improvements with nonlinear over linear decoding. Therefore, one can conclude that the improvements in decoding for midget and OFF units via nonlinear methods can both be primarily attributed to the OFF midget sub-population, which are also better encoders of high-pass details than their parasol counterparts. Previous studies have indeed indicated that midget units may encode more high frequency visual information and that OFF midget units, in particular, exhibit nonlinear encoding properties [12, 30, 31].

**Fig. 8.**
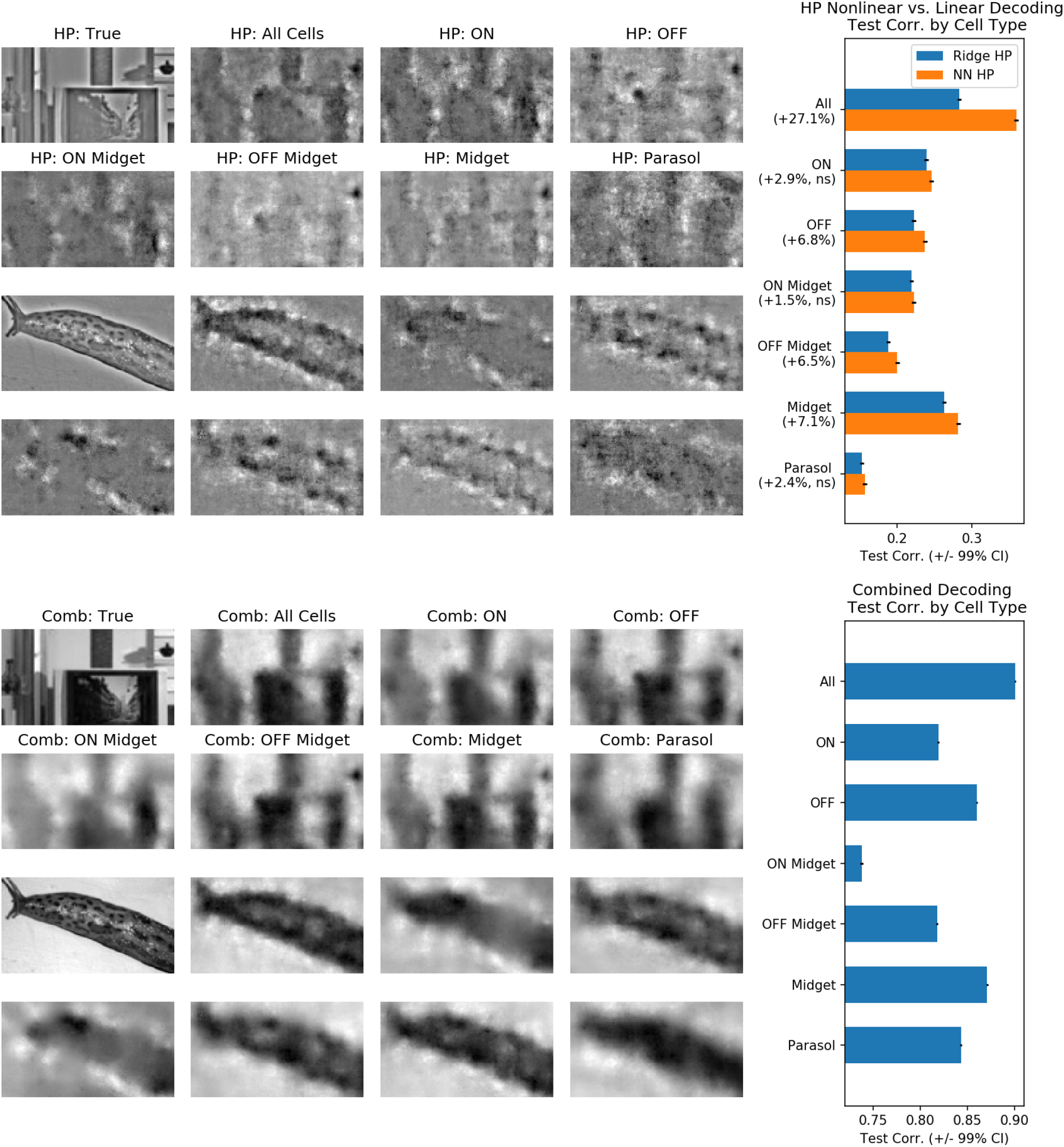
Midget and OFF units contribute most to high-pass, nonlinear decoding. (**Top Left**) Representative high-pass images with their corresponding nonlinear decoded versions using all, ON, OFF, ON Midget, OFF midget, midget, parasol units, respectively. (**Top Right**) Comparison of test correlations between linear and nonlinear high-pass decoding versus cell type. (**Bottom Left**) Representative true images with their corresponding combined decoder outputs using all, ON, OFF, ON Midget, OFF midget, midget, parasol units, respectively. (**Bottom Right**) Comparison of test correlations for the combined decoded images per cell type. Nonlinear decoding most significantly improves midget and OFF cell high-pass and combined decoding while it does not bring any significant benefit to parasol and ON cell decoding.

### A final “deblurring” neural network further improves accuracy, but only in conjunction with nonlinear high-pass decoding

Despite the success of the neural network decoder in extracting more spatial detail than the linear decoder, the combined decoder output still exhibited the blurriness near edges that is characteristic of low-pass image decoding. Therefore we trained a final convolutional “deblurring” network and found that this network was indeed qualitatively able to “sharpen” object edges present in the decoder output images (**Figure 9, Top**; see [22] for a related approach applied to simulated data). Quantitatively, the test pixel-wise correlation improved from 0.890 (±0.0006) and 0.901 (±0.0006) in the linear and combined decoder images, respectively, to 0.912 (±0.0006) in the combined-deblurred images (**Figure 9, Middle; Table 1**). Comparison by SSIM, a more perceptually oriented measure [32], also revealed similar advantages in deblurring in combination with nonlinear decoding over other methods (**Figure 9, Bottom**). In short, this final addition to the decoding scheme brought both subjective and objective improvements to the quality of the final decoder outputs.

**Fig. 9.**
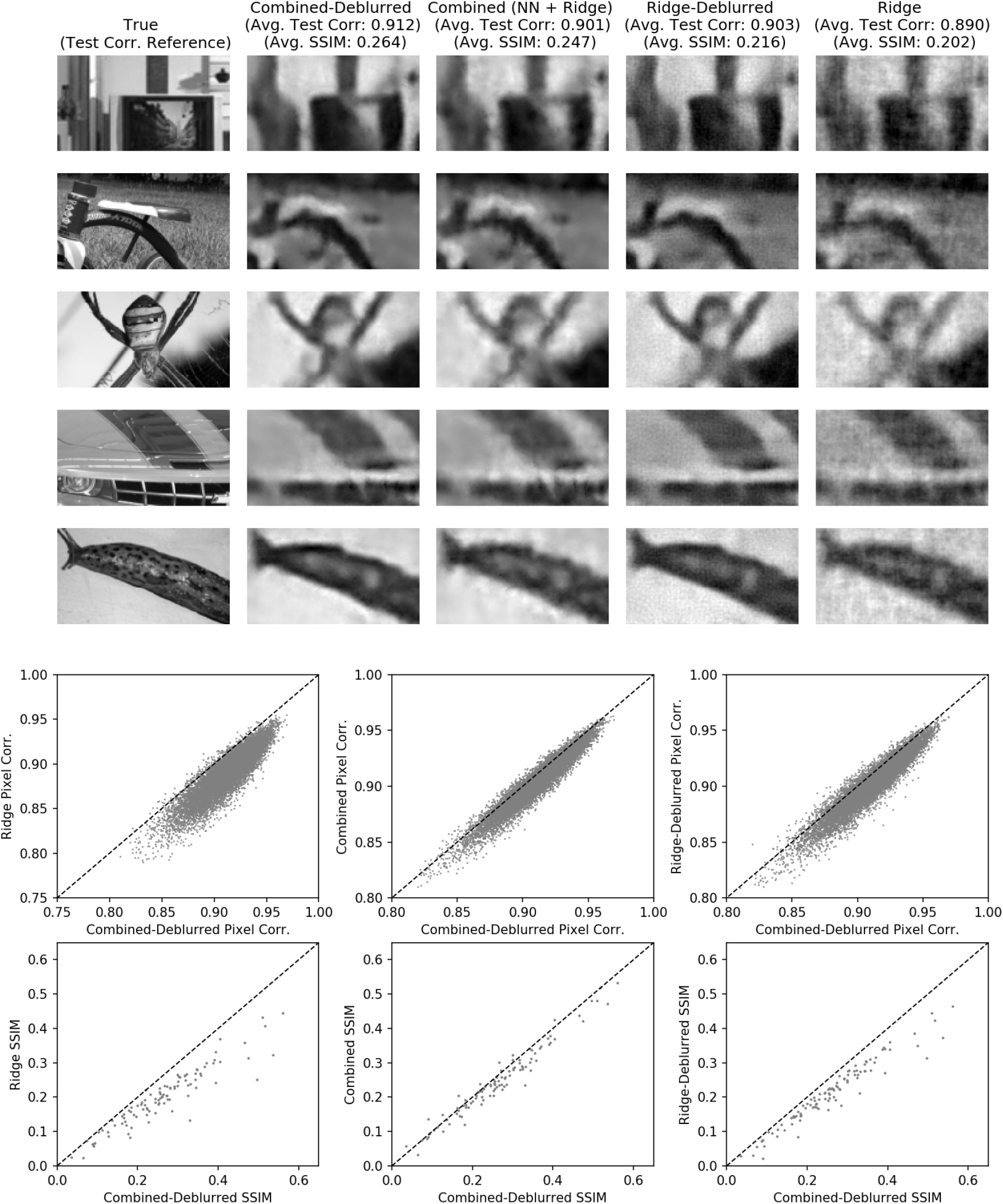
Neural network deblurring further improves nonlinear decoding quality. (**Top**) Representative true images and their corresponding combined-deblurred, combined, ridge-deblurred, and ridge decoder outputs. (**Middle**) Comparisons of pixel-wise test correlation of the combined-deblurred versus ridge, combined, and ridge-deblurred decoder outputs, respectively. (**Bottom**) Comparisons of SSIM values of the combined-deblurred versus ridge, combined, and ridge-deblurred decoder outputs, respectively. The combined-deblurred images had the highest mean SSIM at 0.265 (±0.018, 90% CI) compared to 0.247 (±0.017, 90% CI) and 0.202 (±0.014, 90% CI) for the combined and ridge decoder images, respectively. The ridge-deblurred images had a SSIM of 0.216 (±0.015), which is lower than those of both the combined and the combined-deblurred images. The deblurring network, specifically in combination with nonlinear decoding, brings quantitative and qualitative improvements to the decoded images. See Figure 10 for a similar analysis on a second dataset.

The deblurring network is trained to map noisy, blurry decoded images back to the original true natural image — and therefore implicitly takes advantage of statistical regularities in natural images. (See [22] for further discussion on this point.) Hypothetically, applying the deblurring network to linear decoder outputs could be sufficient for improved decoding. We therefore investigated the necessity of nonlinear decoding in the context of the deblurring network. Re-training and applying the deblurring network on the simple ridge decoder outputs (with the result denoted “ridge-deblurred” images) produced a final mean pixel-wise test correlation of 0.903 (±0.0006), which is lower than that of the combined-deblurred images (**Figure 9; Table 1**). Comparison by SSIM also yielded identical findings. Therefore, enforcing natural image priors on the decoder outputs was largely successful only when the outputs were obtained via nonlinear decoding with minimal noise, demonstrating the necessity of nonlinear decoding within the decoding algorithm.

## Discussion

The approach presented above combines recent innovations in image restoration with prior knowledge of neuronal receptive fields to yield a decoder that is both more accurate and scalable than the previous state of the art. A comparison of linear and nonlinear decoding reveals that linear methods are just as effective as nonlinear approaches for low-pass decoding, while nonlinear methods are necessary for accurate decoding of high-pass image details. The nonlinear decoder was able to take advantage of spike temporal correlations in high-pass decoding while the linear decoder was not; both decoders utilized temporal correlations in low-pass decoding. Furthermore, much of the advantage that nonlinear decoding brings can be attributed to the fact that OFF midget units best encode high-pass visual details in a manner that is more nonlinear than the other RGC types, which aligns with previous findings about the nonlinear encoding properties of this RGC sub-class [31].

These results differ from previous findings (using non-natural stimuli) that linear decoders are unaffected by spike temporal correlations [25, 28] as, evidently, the low-pass linear decoder is just as reliant on such correlations as the nonlinear decoder for low-pass decoding. On the other hand, they also seem to support prior work indicating that nonlinear decoders are able to extract temporally coded information that linear decoders cannot [28, 29]. Indeed, previous studies have noted that retinal cells can encode some characteristics of visual stimuli linearly and others nonlinearly [28, 33–35], which corresponds with our findings that temporally encoded low-pass stimuli information can be recovered linearly while temporally encoded high-pass information cannot. The above may help explain why linear and neural network decoders perform equally well for low-pass images but exhibit significantly different efficacies for high-pass details.

Nevertheless, several key questions remain. While our nonlinear decoder demonstrated state-of-the-art performance in decoding the high-pass images, the neural networks still missed many spatial details from the true image. Although it is unclear how much of these missing details can be theoretically be decoded from spikes from the peripheral retina, we suspect that improvements in nonlinear decoding methods are possible. For example, while our spatially-restricted parameterization of the nonlinear decoder allowed for efficient decoding, it could lose important information in the dimensionality reduction process.

Likewise, the deblurring of the combined decoder outputs is a challenging problem that current image restoration methods in computer vision likely cannot fully capture. Specifically, this step represents an unknown combination of super-resolution, deblurring, denoising, and inpainting. With ongoing advances in image restoration networks that can handle more complex blur kernels and noise, it is likely that further improvements in performance are possible [36–44].

Finally, while our decoding approach helped shed some light on the importance of nonlinear spike temporal correlations and OFF midget cell signals on accurate, high-pass decoding, the specific mechanisms of visual decoding have yet to be fully investigated. Indeed, many other sources of nonlinearity, including nonlinear spatial interactions within RGCs or nonlinear interactions between RGCs or RGC types, are all factors that could help justify nonlinear decoding that we did not explore [33–35, 45–48]. For example, it has been suggested that nonlinear interactions between jointly activated, neighboring ON and OFF cells may signal edges in natural scenes [12]. We hope to investigate these issues further in future work.

## Materials and methods

### RGC datasets

See [12] for full experimental procedures. Briefly, retinae were obtained from terminally anesthetized macaques used by other researchers in accordance with animal ethics guidelines (see **Ethics Statement**). After the eyes were enucleated, only the eye cup was placed in a bicarbonate-buffered Ames’ solution. In a dark setting, retinal patches, roughly 3mm in diameter, were placed with the RGC side facing down on a planar array of 512 extracellular micro-electrodes covering a 1.8mm-by-0.9mm region. For the duration of the recording, the *ex vivo* preparation was perfused with Ames’ solution (30-34 C, pH 7.4) bubbled with 95% O_2_, 5% CO_2_ and the raw voltage traces were bandpass filtered, amplified, and digitized at 20kHz [30, 49–51].

In total, 10,000 natural scene images were displayed with each image being displayed for 100 ms before and after 400 ms intervals of a blank, gray screen. 9,900 images were chosen for training and the remaining 100 for testing. The recorded neural spikes were spike-sorted using the YASS spike sorter to obtain the spiking activities of 2094 RGC units [26], which is significantly more units than previous decoders were trained to decode [12, 23–25]. Due to spike sorting errors, some of these 2094 units may be either over-split (partial-cell) or over-merged (multi-cell). Nevertheless, over-split and over-merged units can still provide decoding information [52] and we therefore chose to include all spike sorted units in the analyses here, in an effort to maximize decoding accuracy. In the LASSO regression analysis (described below), we do perform feature selection to choose the most informative subset of units, reducing the selected population roughly by a factor of two. Finally, to incorporate temporal spike train information, the binary spike responses were time-binned into 10 ms bins (50 bins per displayed image). A second retinal dataset prepared in an identical manner was used to validate our decoding method and accompanying findings (**Supporting Information S1**).

While the displayed images were 160-by-256 in pixel dimensions, we restricted the images to a center portion of size 80-by-144 that corresponded to the placement of the multi-electrode array. To facilitate low-pass and high-pass decoding, each of the train and test images were blurred with a Gaussian blur of *σ* = 4 pixels and radius 3σ to produce the low-pass images. The filter size approximates the average size of the midget RGC. The high-pass images were subsequently produced by subtracting the low-pass images from their corresponding whole images.

### RGC unit matching and classification

To begin with, we obtained spatio-temporal spike-triggered averages (STAs) of the RGC units from their responses to a separate white noise stimulus movie and classified them based on their relative spatial receptive field sizes and the first principal component of their temporal STAs [30]. Afterwards, both MSE and cosine similarity between electrical spike waveforms were used to identify each white noise RGC unit’s best natural scene unit match and vice versa. Specifically, for each identified white noise unit, we chose the natural scene unit with the closest electrical spike waveform using both measures and only kept the white noise units that had the same top natural scene candidate found by both metrics. Then, we performed the same procedure on all natural scene units, keeping only the units that had the same top white noise match using both metrics. Finally, we only kept the white noise-natural scene RGC unit pairs where each member of the pair chose each other as the top match via both MSE and cosine similarity. This ensured one-to-one matching and that no white noise or natural scene RGC was represented more than once in the final matched pairs. In total, 1033 RGC units were matched in this one-to-one fashion, within which there were 72 ON parasol, 87 OFF parasol, 175 ON midget, and 195 OFF midget units. Several other cell types, such as small bistratified and ON/OFF large RGC units, were also found in smaller numbers. We also confirmed that the top 25 units chosen per pixel by LASSO, which comprise the 805 unique units feeding into the nonlinear decoder, also represented the four main RGC classes proportionally.

We chose a very conservative matching strategy to ensure one-to-one representation and maximize the confidence in the classification of the natural scene units. Naturally, such a matching scheme produced many unmatched natural scene units and a smaller number of unmatched white noise units. On average, the unmatched natural scene units had similar firing rates to the matched units while having smaller maximum channel spike waveform peak-to-peak magnitudes. While it is likely that a relaxation of matching requirements would yield more matched pairs, we confirmed that our matching strategy still resulted in full coverage of the stimulus area by each of the four RGC types (**Supporting Information S2**).

### Low-pass linear decoding

To perform efficient linear decoding on a large neural spike matrix without over-fitting, for each RGC, we summed spikes within the 30 - 150 ms and 170 - 300 ms time bins, which correspond to the image onset and offset response windows. Thus, with *n, t, x* indexing the RGC units, training images, and pixels, respectively, the RGC spikes were organized into matrix 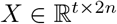 and the training images into matrix 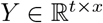. To initially solve the linear equation *Y = Xβ*, the weights were inferred through the expression 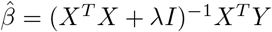, in which the regularization parameter λ ≈ 4833 was selected via three-fold cross-validation on the training set [27]. Although we reduced the number of per-image time bins from 50 to 2, we confirmed that performing ridge regression on the augmented 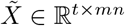 with *m* indexing the 50 time bins yielded essentially identical low-pass decoding performance, as discussed in the Results section.

Additionally, to perform pixel-specific feature selection for high-pass decoding, we performed LASSO regression [27], which was proven to successfully select for relevant units, on the same neural bin matrix *X* from above [25]. Due to the enormity of the neural bin matrix, *Celer,* a recently developed accelerated L1 solver, was utilized to individually set each pixel’s L1 regularization parameter as decoding each pixel represents an independent regression sub-task [53].

### High-pass nonlinear decoding

To maximize high-pass decoding efficacy with the nonlinear decoder, the augmented 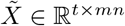 was chosen as the training neural bin matrix. As noted above, nonlinear methods, including kernel ridge regression and feedforward neural networks, have been successfully applied to decode both the locations of black disks on white backgrounds [25] and natural scene images [23]. Notably the former study utilized L1 sparsification of the neural response matrix so that only a handful of RGC responses contributed to each pixel before applying kernel ridge regression. We borrow this idea of using L1 regression to create a sparse mapping between RGC units and pixels before applying our own neural network decoding as explained below. However, the successful applications of feedforward decoding networks above crucially depended on the fact that they utilized a small number of RGCs (91 RGCs with 5460 input values and 90 RGCs with 90 input values, respectively). For reference, constructing a feedforward network for our spike data of 2094 RGC units and 104,700 inputs would yield an infeasibly large number of parameters in the first feedforward layer alone. Similarly, kernel ridge regression, which is more time-consuming than a feedforward network, would be even more impractical for large neural datasets.

Therefore we constructed a spatially-restricted network based on the fact that each RGC’s receptive field encodes a small subset of the pixels and, conversely, each pixel is represented by a small number of RGCs. Specifically, each unit’s image-specific response *m*-vector is featurized to a reduced *f*-vector so that each unit is assigned its own featurization mapping that is preserved across all pixels. Afterwards, for each pixel, the featurized response vectors of the *k* most relevant units are gathered into a *fk*-vector and further processed by nonlinear layers to produce a final pixel intensity value. The *k* relevant units are derived from the L1 weight matrix 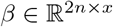 from above. Within each pixel’s weight vector 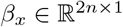 and an individual unit’s pixel-specific weights 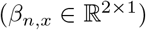, we calculate the L1-norm λ_*x,n*_ = |*β_n,x_*|_1_ and select the units corresponding to the *k* largest norms for each pixel. The resulting high-pass decoded images are added to the low-pass decoded images to produce the combined decoder output. Note that while the RGC featurization weights are shared across all pixels, each pixel has its own optimized set of nonlinear decoding weights (**Figure 2**).

The hyperparameters *f* = 5, *k* = 25 were chosen from an exhuastive grid search spanning *f* ∈ {5, 10, 15, 20}, *k* ∈ {5, 10, 15, 20, 25} so that the values at which no further performance gains were observed were selected. The neural network itself was trained with a variant of the traditional stochastic gradient descent (SGD) optimizer that includes a momentum term to speed up training [54] (momentum hyperparameter of 0.9, learning rate of 0.1, and weight regularization of 5.0 * 10^-6^ used for training the network over 32 epochs).

### Deblurring network

To further improve the quality of the decoded images, we sought to borrow image restoration techniques from the ever-growing domain of neural network-based deblurring. Specifically, a deblurring network leveraging natural image priors would take in the combined decoder outputs and produce sharpened versions of the inputs. However, these networks usually come with high requirements for training dataset size and only using the 100 decoded images corresponding to the originally held out test images would be insufficient.

As a result, we sought to virtually augment our decoder training dataset of 9,900 spikes-image pairs for use as training examples in the deblurring scheme. The 9,900 training spikes-image pairs were sub-divided into ten subsets of 990 pairs. Then, each subset was held out and decoded (both linearly and nonlinearly) with the other nine subsets used as the decoders’ training examples. Rotating and repeating through each of the ten subsets allowed for all 9,900 training examples to be transformed into test-quality decoder outputs, which could be used to train the deblurring network. (To be clear, 100 of the original 10,000 spikes-images pairs were held out for final evaluation of the deblurring network, with no data leakage between these 100 test pairs and the 9,900 training pairs obtained through the above dataset augmentation.) An existing alternative method would be to craft and utilize a generative model for artificial neural spikes corresponding to any arbitrary input image [22, 23]. However, the search for a solution for the encoding problem is still a topic of active investigation in neuroscience; our method circumvents this need for a forward generative model.

With a sufficiently large set of decoder outputs, we could adopt well-established neural network methods for image deblurring and super-resolution [36–44]. Specifically, we chose the convolutional generator of DeblurGANv2, an improvement of the widely adopted DeblurGAN with superior deblurring capabilities [39]. After performing a grid search of the generator ResNet block number hyperparameter ranging {1, 2,…, 7, 8}, the 6-block generator was chosen for training under the Adam optimizer [55] for 32 epochs at an initial learning rate of 1 × 10^-5^ that was reduced by half every 8 epochs.

We do not expect that the decoded images will be near-perfect replicas of the original image. Recordings here were taken from the peripheral retina, where spatial acuity is lower; as a result, one would expect the neural decoding of the stimuli to miss some of the fine details of the original image. Therefore, while the original DeblurGANv2 paper includes pixel-wise L1 loss, a VGG discriminator-based content/perceptual loss, and an additional adversarial loss during training, we excluded the final adversarial loss term, due to the fact that the deblurred images of the decoder would not be perfect (or near-perfect) look-alikes of the raw stimuli images. Instead, we focus on improving the perceptual qualities of the output image, including edge sharpness and contrast, for more facile visual identification. We use both pixel-wise L1 loss and L1 loss between the features extracted from the true images and from the reconstructions in the 3rd convolutional layer of the pre-trained VGG-19 network before the corresponding pooling layer [38, 56].

### Ethics Statement

Eyes were removed from terminally anesthetized macaque monkeys (Macaca mulatta, Macaca fascicularis) used by other laboratories in the course of their experiments, in accordance with the Institutional Animal Care and Use Committee guidelines. All of the animals were handled according to approved institutional animal care and use committee (IACUC) protocols (28860) of the Stanford University. The protocol was approved by the Administrative Panel on Laboratory Animal Care of the Stanford University (Assurance Number: A3213-01).

## Supporting information

### S1 Fig. Validation of decoding methods on second RGC dataset

**Fig 10.**
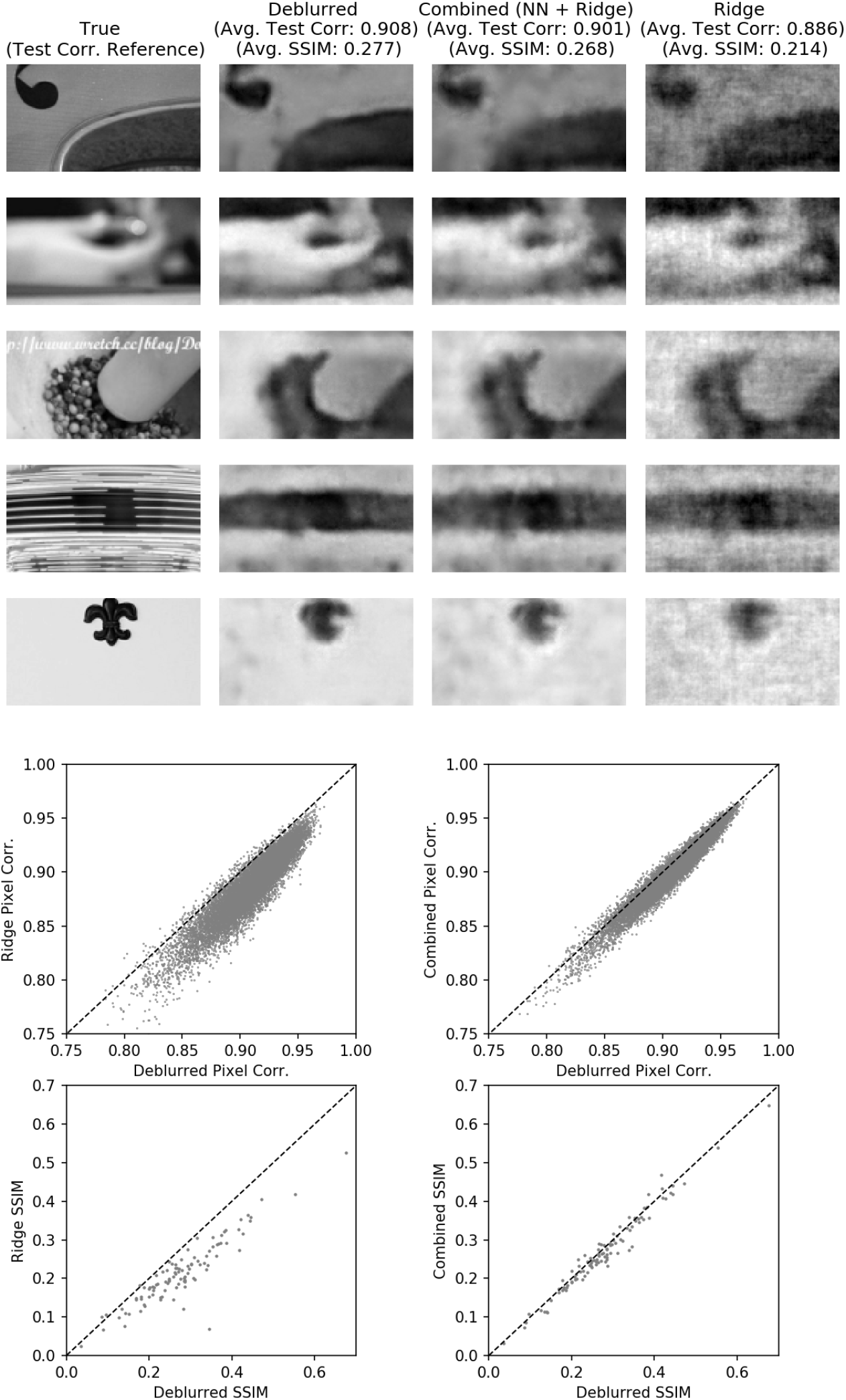
Decoding method results corroborated on second RGC dataset. (**Top**) Representative outputs from the decoding algorithm compared to those from a simple linear decoder. (**Bottom**) Comparison of pixel-wise test correlations and SSIM values between deblurred and linear decoder outputs and against combined decoder outputs, respectively. The second dataset consisted of the responses of 1987 RGC units to 10,000 images, prepared in an identical manner as the first dataset. The superiority of nonlinear decoding with deblurring is apparent.

### S2 Fig. Matching of white noise and natural scene RGC units

Because hundreds of white noise and more than a thousand natural scene RGC units were discarded during the matching process, these unmatched units were analyzed to see whether they exhibited any distinguishing properties from the matched units. Comparing the mean firing rates of the matched and unmatched units revealed no clear differences: 10.53 Hz vs. 11.46 Hz for matched and unmatched natural scene units; 6.56 Hz vs. 7.03 Hz for matched and unmatched white noise units. However, the mean maximum channel peak-to-peak values (PTPs) were markedly different between matched and unmatched units within both experimental settings: 22.06 vs. 10.21 for matched and unmatched natural scene units; 24.93 vs. 18.48 for matched and unmatched white noise units.

Non-matching of units is likely caused by several factors. To begin with, MSE and cosine similarity are not perfect measures of template similarity. Many close candidates were quite similar in shape to the reference templates, but either had a slightly different amplitude or had peaks and troughs at different temporal locations. Indeed it is possible that using a more flexible similarity metric would recover more matching units. Meanwhile, it is also likely that some of the unmatched units in either experimental setting are simply inactive units. Specifically, it could be the case that some units are inactive during white noise stimulation, but more active for natural scene input and vice versa. Finally, difficulties with spike sorting smaller units could also lead to mismatches. Nevertheless, despite the above issues, we were able to recover full coverage of the stimulus region for each cell type, as shown in **Figure 11**.

**Fig 11.**
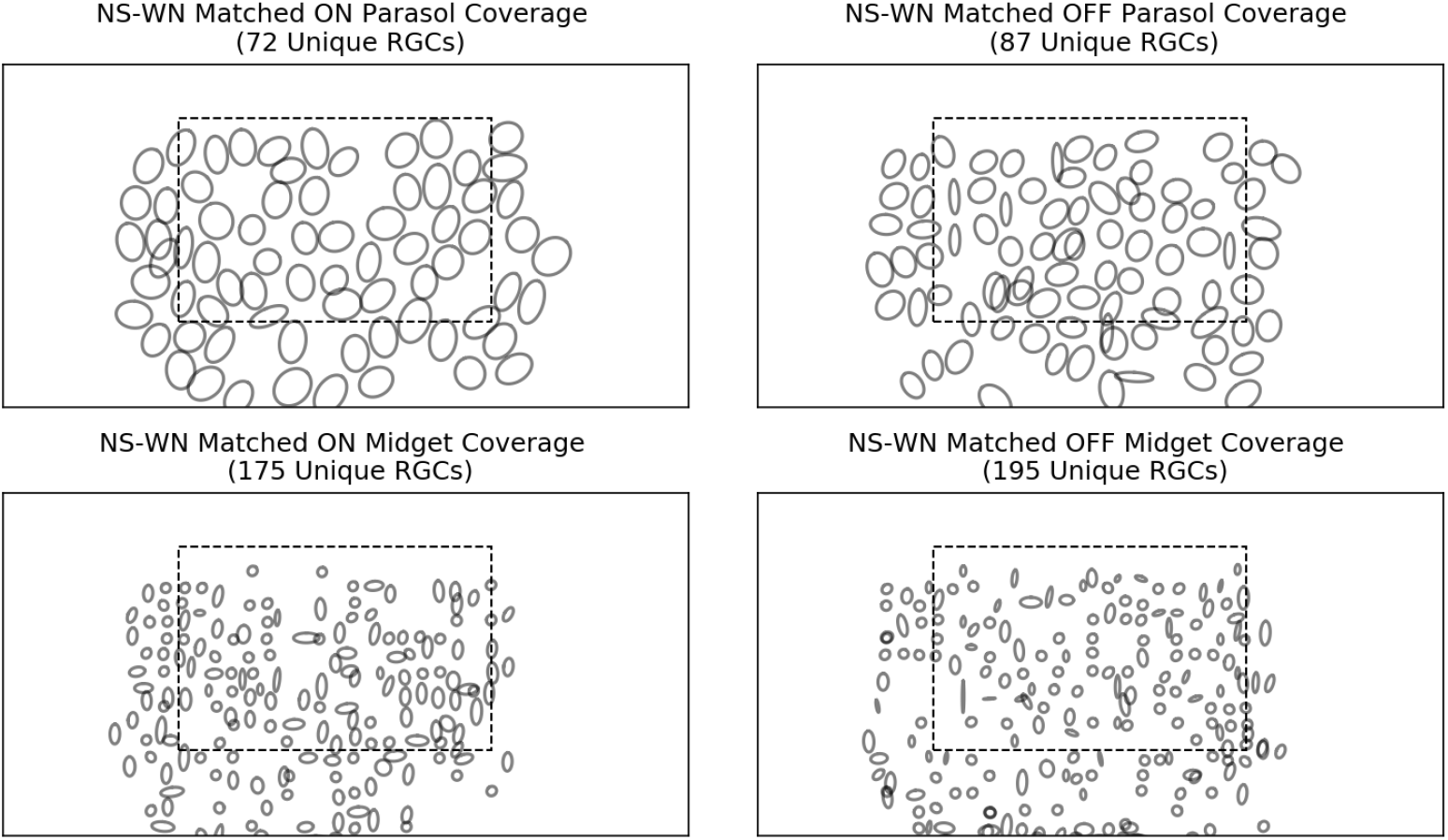
Coverage of image area by matched RGC cells. All four cell types, ON/OFF parasol/midget, sufficiently cover the image area (marked in dashed rectangle) with the receptive fields of their constituent white noise-natural scene matched units.

## Acknowledgments

We thank Eric Wu and Nishal Shah for helpful discussions.

## Footnotes

1 Note that these correlation values are much higher than the subsequent correlation values in this manuscript as these low-pass decoded images were evaluated against the true low-pass images, which are much easier decoding targets than the true whole images themselves.

